# Modeling flexible protein structure with AlphaFold2 and cross-linking mass spectrometry

**DOI:** 10.1101/2023.09.11.557128

**Authors:** Karen Manalastas-Cantos, Kish R. Adoni, Matthias Pfeifer, Birgit Märtens, Kay Grünewald, Konstantinos Thalassinos, Maya Topf

## Abstract

We propose a pipeline that combines AlphaFold2 (AF2) and crosslinking mass spectrometry (XL-MS) to model the structure of proteins with multiple conformations. The pipeline consists of two main steps: ensemble generation using AF2, and conformer selection using XL-MS data. For conformer selection, we developed two scores – the monolink probability score (MP) and the crosslink probability score (XLP), both of which are based on residue depth. We benchmarked MP and XLP on a large dataset of decoy protein structures, and showed that our scores outperform previously developed scores. We then tested our methodology on three proteins having an open and closed conformation in the Protein Data Bank: Complement component 3 (C3), luciferase, and glutamine-binding periplasmic protein (QBP), first generating ensembles using AF2, which were then screened for the open and closed conformations using experimental XL-MS data. In five out of six cases, the most accurate model within the AF2 ensembles – or a conformation within 1 Å of this model – was identified using crosslinks, as assessed through the XLP score. In the remaining case, only the monolinks (assessed through the MP score) successfully identified the open conformation of QBP. This serves as a compelling proof-of-concept for the effectiveness of monolinks. In contrast, the AF2 assessment score (pTM) was only able to identify the most accurate conformation in two out of six cases. Our results highlight the complementarity of AF2 with experimental methods like XL-MS, with the MP and XLP scores providing reliable metrics to assess the quality of the predicted models.

## Introduction

AlphaFold2 (AF2) has revolutionized structural biology with unprecedented accuracy in protein structure prediction, even on sequences for which related structures are unavailable (1). AF2 has been used to exhaustively predict the structures of more than 200 million UniProt sequences (2), which is both an invaluable resource to the structural biology community, as well as a test of AF2’s current structure prediction capacity. In particular, it has been shown that AF2 predicts ordered protein domain structures well, but performs less well on proteins with predicted flexibility or disorder (3). A possible cause is that AF2 and other protein structure prediction approaches have been trained with the assumption that one protein sequence corresponds to one structure, something we know to be untrue for a wide variety of proteins, such as molecular switches which transition between two different conformations as part of their function (4), as well as proteins that are either fully or partially disordered. The issue is compounded by the fact that the Protein Data Bank (PDB) (5)—the protein structure database used to train structure prediction models—is biased towards proteins that are relatively ordered, since disordered regions are usually recalcitrant to X-ray crystallography, and are thus poorly-represented (6). In addition, proteins that have multiple possible states resulting from domain rearrangements (e.g. upon binding to other proteins) may not have all of their conformations documented in the PDB.

To its credit, AF2 outputs error estimates for its structure predictions, and usually assigns low confidence to intrinsically-disordered regions of a protein. In the case of molecular switches that have relatively-rigid domains that undergo inter-domain movements, AF2’s predicted aligned error (PAE) is sometimes predictive of boundaries between domains, but not always (Fig. 1A).

**Fig. 1.**
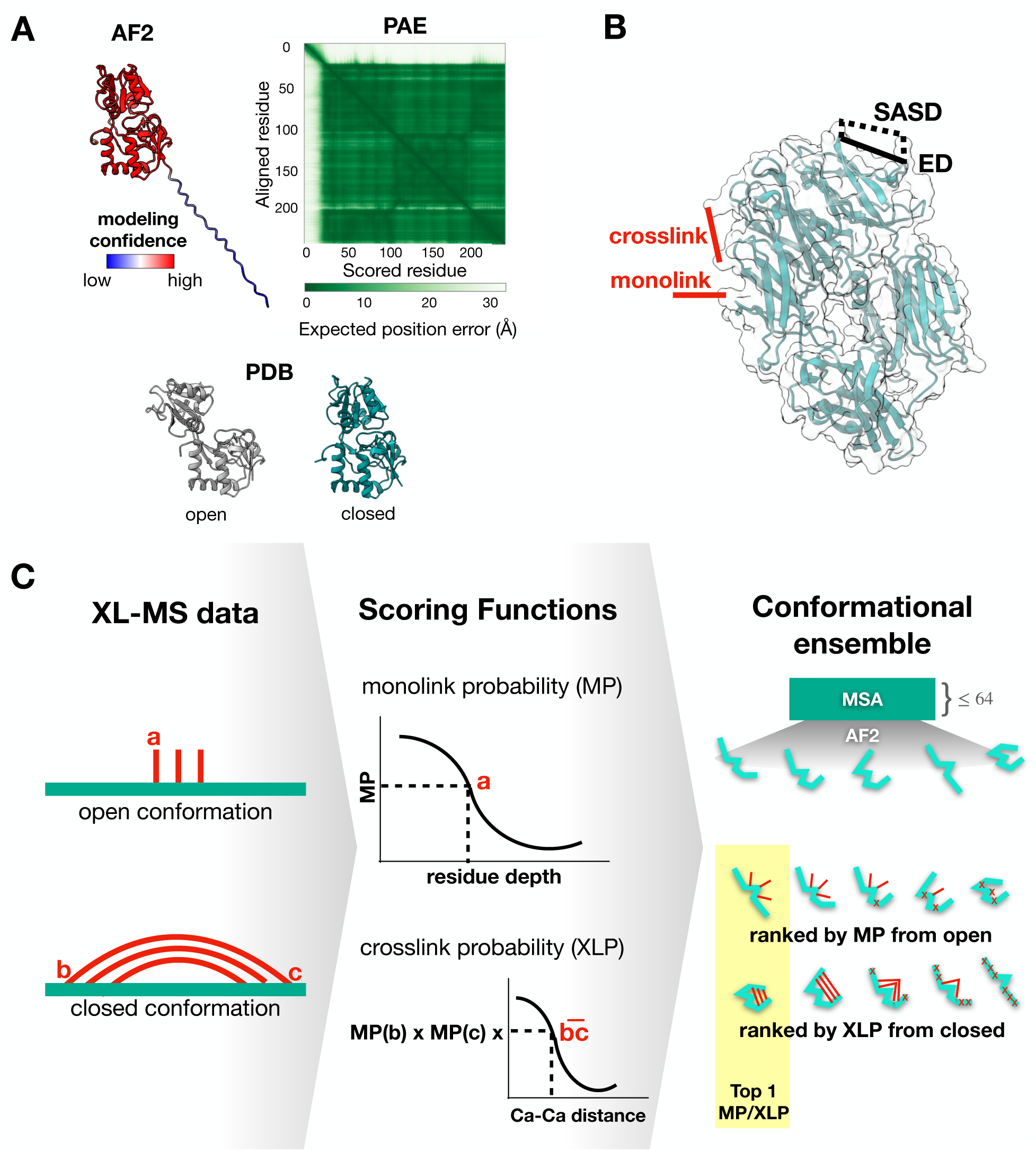
(A) AlphaFold2 default prediction and the predicted aligned error (PAE) for glutamine-binding periplasmic protein (QBP), along with the PDB structures for QBP open and closed conformations. AF2 predicted only the closed conformation of QBP with high confidence, and was not able to recognize the flexibility between the two domains. Blocks of high PAE (white regions on plot) typically indicate boundaries between domains, but QBP was predicted as a single, low PAE block (high PAE region is caused by the unfolded N-terminus). (B) Crosslinks occur when both ends of a chemical crosslinker (red) bond covalently with the protein, while monolinks involve only one attachment site. The Euclidean distance (ED) is the shortest path between the crosslinked C⍺ atoms, while the solvent-accessible surface distance (SASD) is the shortest path between the crosslinked C⍺ atoms along the surface of the protein. (C) Our approach for flexible protein modeling involves acquiring XL-MS data in the form of monolinks (a) and crosslinks (bc) from the open and closed conformations, while predicting an ensemble of models using AlphaFold2 with shallow multiple sequence alignments (≤ 64 sequences). The AF2 models are scored by how well they match XL-MS data from each conformation, using the monolink (MP) and crosslink probability (XLP) scores described in Experimental Procedures.

Thus, experimental methods that can capture protein dynamics remain critical in a post-AF2 world. For example, in cross-linking mass-spectrometry (XL-MS), proteins are incubated at near-physiological conditions with chemical crosslinkers such as the most commonly-used reagents disuccinimidyl suberate (DSS) and bissulfosuccinimidyl suberate (BS3), which both bind with highest affinity to lysine residues on the protein surface (7). The linker may attach at only one end, giving rise to a monolink, or both ends of the same linker may attach to different parts of the protein, giving rise to a crosslink (Fig. 1B). Analysis of the linker attachment sites by MS gives information about what parts of the protein are on the surface. Additionally, crosslinks can constrain the maximum distance between the two sites on the protein where they are attached, since the linker has a defined length (11.4Å for DSS and BS3) (8). Crosslinks have been used to evaluate protein structure models by either flagging maximum distance violations, where the maximum C⍺-C⍺ Euclidean distance that can be spanned by the linker is exceeded (9–12), or by more sophisticated scoring functions that are based on the solvent-accessible surface distance (SASD) (Fig. 1B). The SASD, while more time-consuming to compute, is a better approximation of physical conditions, and was shown to perform better than ED in distinguishing near-native protein structures from a group of decoys (13–17).

The primary tradeoff of methods like XL-MS that can be performed in solution is that they typically do not give atomic resolution information. For this reason, a common experimental pipeline when studying flexible proteins is to combine both high-resolution structural information from X-ray crystallography, for example, with measurements in solution such as small-angle scattering to quantify flexibility, and XL-MS or FRET to give distance constraints that might not be present in the crystal conformation. In the absence of experimentally-determined structures, homology-based models of proteins have also been used in combination with distance constraints measured in solution in order to model the structure of flexible proteins. However, with the advent of deep learning-based protein structure prediction methods such as AF2, this pipeline could be performed on proteins that have no sequence homology to structures in the PDB, or proteins for which only some of the conformations are known.

In this paper, we present how XL-MS data can be used to supplement structure predictions from AlphaFold2, particularly on flexible proteins (Fig. 1C). It has previously been shown that AF2 generates a wider variety of conformations when using a shallow multiple-sequence alignment (MSA), than when not limiting alignment depth (18). We leverage this observation to generate an ensemble of conformations for a set of flexible proteins that have an open and a closed conformation deposited in the PDB. We screened these AF2 predictions using experimental XL-MS data using two scoring functions we have developed: the crosslink and monolink probability scores, XLP and MP respectively. We show that XLP and MP are able to select for the open and closed conformations from the AF2 ensembles. Furthermore, we present a case in which monolinks are more informative than crosslinks in selecting the experimentally-relevant conformer from an ensemble. Thus, we propose a general pipeline to model flexible proteins by first generating an ensemble with AF2, then identifying the conformer present in a given experimental condition using crosslinks and monolinks from XL-MS.

## Experimental Procedures

### Defining scoring functions using XL-MS data

We defined two new scoring functions to distinguish near-native structures from a set of decoy protein structures, based on distributions of crosslinks and monolinks obtained from XL-MS experiments. Experimentally-determined BS3 and DSS crosslinks and monolinks that were mapped to PDB structures were first obtained from the XlinkAnalyzer database (https://www.embl-hamburg.de/XlinkAnalyzer/database.html) (19). A total of 831 unique lysine residues across 43 PDB structures were found to be tagged with either BS3 or DSS. The depths of both tagged and untagged lysines were computed in the respective PDB structures using the program EDTSurf, which is a fast algorithm for generating protein surfaces, as well as calculating depths to this surface, using Euclidean distance transform (20, 21). The following parameters were used to run EDTSurf: probe radius=2.5, scale=0.5.

The residue depths computed by EDTSurf were compared to those obtained from DEPTH, which is a more accurate but slower method that places the protein structure in a random orientation in a box of explicit solvent for several iterations, while computing the distance of each residue from the nearest bulk solvent (22). Using the XlinkAnalyzer dataset, we show that the faster EDTSurf was able to approximate DEPTH to a Pearson cross-correlation coefficient of 0.97 (Fig. S1). Thus, unless otherwise stated, residue depths mentioned in this work were computed with the Euclidean distance transform algorithm (EDT).

The depth distribution of tagged lysines is shown in Fig. 2A, along with the depth distribution of all lysines across the 43 PDB structures assessed. In addition, the Euclidean distances between C⍺ atoms of crosslinked lysines were computed, with the distribution of distances shown in Fig. 2B. The depth distributions were fitted with a negative power function, while the C⍺-C⍺ distance distribution was fitted with a sigmoidal function using SciPy v1.5.2 (23) (Fig. 2).

**Fig. 2.**
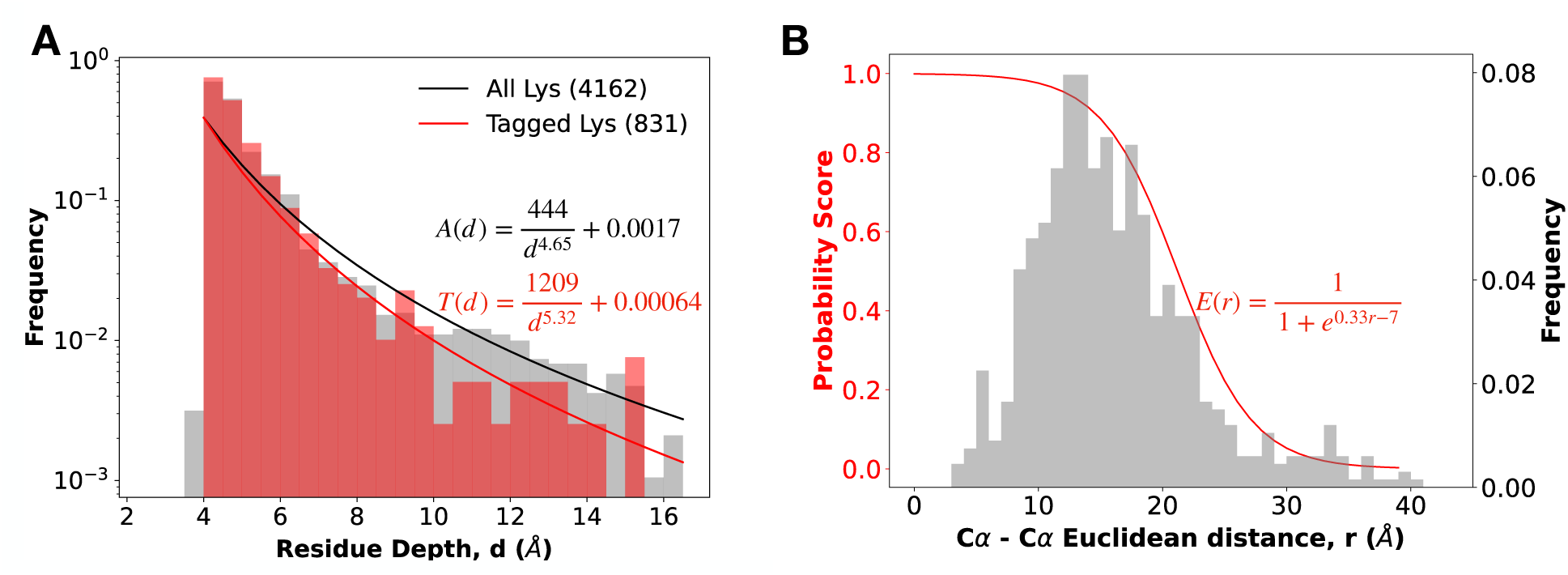
Empirical distributions of (A) depths and (B) C⍺-C⍺ Euclidean distances of DSS- and BS3-crosslinked lysine residues. 831 unique lysine residues were found with an attached linker out of 4162 lysines across 43 PDB structures. Depth distributions of tagged and untagged lysines were fit with a negative power function, while the C⍺-C⍺ distance distribution was fit with a sigmoidal function as shown.

The monolink probability (MP) score for a monolink in a given structure is thus defined as follows:

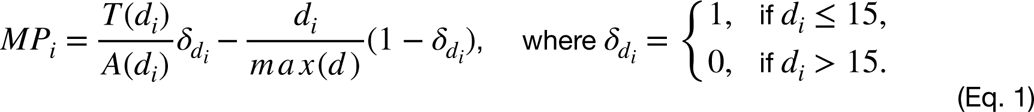

with the first term corresponding to the ratio between tag probability *T* and lysine occurrence *A* at a given depth *d_i_* in the structure (Fig. 2A), and the second term being a penalty for monolinks that are a depth of more than 15Å in the structure, with the 15Å cutoff set empirically from the observed residue depths. The MP score of a given model is the averaged MP score of all monolinks found in the structure.

The crosslink probability (XLP) score for a crosslink spanning residues and *j* in a given structure is then defined as:

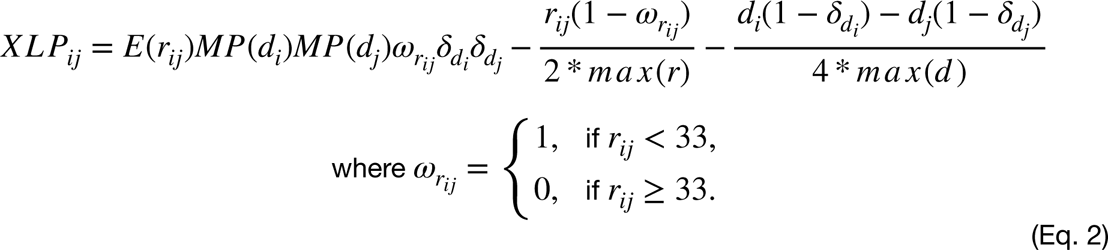

with the first term corresponding to the likelihood *E* of the C⍺-C⍺ distance r_ij_being spanned by a BS3/DSS crosslink (Fig. 2B), weighted by the monolink probability *MP* at both residues and *j*. The second term is a penalty for C⍺-C⍺ distances equal to or exceeding 33Å, while the third term is a penalty for either residue exceeding a maximum depth of 15Å, as in the MP score. A threshold of 33Å was set, as this is the maximum spanning distance for DSS and BS3 previously observed (10, 16). The weights for the penalty terms were set to 0.5 for C⍺-C⍺ maximum distance violations, and 0.25 for maximum residue depth violations. Similar to the MP score, XLP is averaged across all crosslinks found in the structure.

Unlike the monolink (24) and crosslink (16) scores we previously developed, both MP and XLP were formulated to scale smoothly from -1 to 1, with a score of 1 indicating that the given structure provides the highest probability that the given monolink or crosslink can be formed. Having such a defined scale means that the magnitude of the scores of a single model is independent of the distribution of other predicted models, and thus does not depend on the assumption that the models are normally-distributed, nor that the ensemble contains the correct model, unlike in other methods of score normalization, such as the computation of a z-score.

### Benchmarking MP and XLP scores

The MP and XLP scores were evaluated on the structure decoy dataset 3DRobot, a set of 200 nonhomologous proteins randomly-selected from the PDB, each with 300 structural decoys with root-mean-squared deviation (RMSD) from the native structure ranging from 0 to 12Å (25). Sets of “simulated” crosslinks and monolinks were generated at different recovery rates using the native structures in the 3DRobot dataset. Similar to our previous work (24), we defined simulated monolinks as lysine residues in the native structure with a depth of 6.25Å or less, as computed by DEPTH (22). Simulated crosslinks were defined as pairs of lysine residues in the native structure with a depth of at most 6.25Å, and with an C⍺-C⍺ SASD of at most 33Å, computed from the structure with Jwalk (16). We simulated a range of recovery rates (i.e. the fraction of detected crosslinks from all possible crosslinking events) from 10-100%, in increments of 10%. At each recovery rate, all possible crosslinking events were randomly sampled 1000 times, to simulate variation that may occur in XL-MS experiments. Similarly, we generated simulated monolinks in this bootstrapped manner, by randomly sampling all possible monolinks 1000 times at a range of recovery rates of 10-100%.

The XLP and MP scores were calculated for each 3DRobot structure at each recovery rate, averaged across the 1000 samplings, and compared with the previously described MNXL (Matched and Nonaccessible Crosslinks) score.

MNXL is defined as follows:

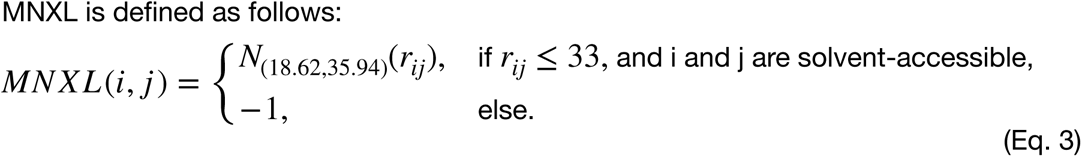

where *i* and *j* are the crosslinked residues. r_ij_ is the SASD between them, and *N*_(18.62,35.94)_ is the normal distribution calculated from all SASDs ≤ 33Å from the crosslink database XLdb (16).

A well-performing scoring function assigns higher ranks to structures that closely resemble the native structure, and can effectively distinguish near-native structures from less accurate decoys. To assess ranking performance, we calculated the Spearman cross-correlation (CC) between the scores and the RMSD from the native structure. In addition, we measured each score’s ability to identify near-native structures by computing the area under the receiver operating characteristic curve (AUC). The receiver operating characteristic curve (ROC) plots the relationship between the number of true and false positives at different scoring thresholds. In our case, true positives were defined as near-native models (TM-score ≥ 0.9) scoring above the threshold, while false positives were non-native models (TM-score < 0.9) also scoring above the threshold (26).

To determine if there were significant performance differences between XLP and MNXL, we compared the Spearman CC and AUC at each recovery rate using the Wilcoxon signed-rank test implemented in SciPy v1.5.2 (23).

### Collating an experimental XL-MS dataset of flexible proteins

We compiled a test dataset of three proteins with experimental BS3 and/or DSS XL-MS data from the literature (Table S1). Each protein in the dataset had at least one “open” and one “closed” conformation in the PDB, which were defined based on their radius of gyration (Rg) (27–29). Details of the dataset are summarized in Table 1. For glutamine-binding periplasmic protein (QBP), no crosslinks or monolinks were reported for the open conformation. Simulating crosslinks and monolinks using our criteria from the previous section, we found that QBP has one lysine that is exposed only in the open conformation. To test whether this lysine is sufficient to distinguish QBP open conformation, QBP was produced and purified for subsequent XL-MS experiments, as described below.

**Table 1.** Flexible proteins with PDB structures and experimental XL-MS data.

| Protein name<br>(UniProt ID) | RMSD between<br>conformations (Å) | PDBID,<br>open | $R_g$, open<br>(Å) | PDBID,<br>closed | $R_g$,<br>closed (Å) | Experimental<br>XL-MS data |
| --- | --- | --- | --- | --- | --- | --- |
| Complement<br>component 3, C3<br>(P01024) | 26.9 | 2i07 | 46.2 | 2a73 | 42.9 | crosslinks<br>(open and<br>closed)<br>(27) |
| Luciferase<br>(P08659) | 5.4 | 1lci | 24.6 | 4g36 | 23.3 | crosslinks<br>(open and<br>closed)<br>(29) |
| Glutamine-binding<br>periplasmic<br>protein, QBP<br>(P0AEQ3) | 5.3 | 1ggg | 19.0 | 1wdn | 17.5 | crosslinks<br>(closed)<br>(28)<br>monolinks<br>(open, this<br>study) |

### Cloning, expression and purification of QBP

To obtain a C-terminal 6xHis-tagged QBP variant for purification, the open reading frame of QBP was amplified from *E.coli* K12 DNA and cloned into bacterial expression vector pET28a-LIC (Addgene #26094). Transformed *E.coli* BL21 (DE3), cells were cultured in Terrific Broth (ROTH, Germany) at 37°C to an OD_600_ = 0.8 followed by Isopropyl β-D-1-thiogalactopyranoside (IPTG, 500µM) induction of QBP expression for 5 h at 30 °C. Subsequently, cells were harvested, resuspended in lysis buffer (50 mM NaH_2_PO_4_; 300 mM NaCl; 10 mM imidazole; pH 8: supplemented with protease inhibitor cocktail (cOmplete, Roche, Switzerland), DNaseI (ITW REAGENTS, Spain), lysozyme (Sigma-Aldrich, US)) and sonicated. QBP was purified from whole cell lysate in a three-step process using an ÄktaGo chromatography system (Cytiva, USA). First, cleared lysate was loaded onto a HisTrapHP column (Cytiva, US) and washed with buffer A (50 mM NaH_2_PO_4_; 300 mM NaCl; 20 mM imidazole; pH 8). QBP was eluted in a buffer B (50 mM NaH_2_PO_4_; 300 mM NaCl; 500 mM imidazole; pH 8). Second, for release of bound glutamine guanidine hydrochloride was added to a final concentration of 6 M. Lastly, QBP was gradually dialysed in size exclusion buffer (20 mM HEPES pH 7.5; 150 mM NaCl) and purified through size exclusion chromatography (Superdex200, Cytiva, US). Samples of *E.coli* cultures were harvested and resuspended in *E.coli* lysis reagent (NEB, US). Lysates and purified proteins were separated by 4-12% Bis-Tris SDS-PAGE (ThermoFisher Scientific, US) and visualized by Quick coomassie stain (SERVA, Germany, Fig. S2).

### Sample crosslinking and preparation for mass spectrometry

QBP was buffer-exchanged into 20 mM HEPES, pH 7.5, at 4°C (repeated for 10 cycles) to remove salt using an Amicon Ultra-0.5 Centrifugal 10 kDa filter unit (Merck), before dilution to 5.00 µM in 20 mM HEPES. For glutamine-bound QBP, 5 mM glutamine was added prior to crosslinking. 50 mM BS3-d_0_ crosslinker was then added to glutamine-bound QBP, while 50 mM BS3-d_4_ crosslinker was added to glutamine-free QBP. Samples were then incubated at room temperature for 120 min before quenching with 20 mM ammonium bicarbonate. The glutamine-bound QBP with BS3-d_0_ and glutamine-free QBP with BS3-d_4_ were pooled before mass spectrometry analysis. Samples were reduced with 8 mM dithiothreitol (DTT) and incubated at 37°C for 45 min before alkylation using 20 mM iodoacetamide with incubation in darkness for 45 min, followed by a 1:5 dilution with 50 mM ammonium bicarbonate. Trypsin (Promega) was then added at a Trypsin:QBP mass ratio of 1:50 and incubated overnight at 37°C. The digestion was then quenched by addition of trifluoroacetic acid to a final concentration of 0.50%. Samples were ZipTip desalted (Merck) before reconstitution in 1.0% formic acid at a concentration of 1.0 μg/μL.

### Liquid Chromatography-Mass Spectrometry setup, data collection and analysis

The UltiMateTM 3000 RSLCnano liquid chromatography system (Thermo Fisher Scientific), with a 50 cm µPAC™ Neo HPLC analytical column (COL-NANO050NEOB) and a 0.075 mm x 20 mm trap cartridge (Acclaim PepMap C18 100 Å, 3 μm), was connected to a SilicaTip emitter. Column temperature was set at 45°C and column flow rate was set to 300 nL/min. Mobile phase A (0.1% formic acid, 3.2% acetonitrile) and mobile phase B (96.8% acetonitrile/0.1% formic acid) were applied with an elution gradient from 5.0 to 35.0% mobile phase B over 20 min. Total run time per sample was 30 min.

The Thermo Fisher Scientific Orbitrap Eclipse Tribrid mass spectrometer was used for this work. The mass spectrometer was externally calibrated using Pierce^TM^ FlexMix^TM^ calibration solution. nanoESI was performed by the application of a voltage to a SilicaTip emitter, via a HPLC liquid junction cross. Spray stability and intensity was optimized by varying the SilicaTip electrospray voltage (1.0 kV – 10 kV) and varying SilicaTip positioning (in the x, y and z dimensions). Transfer capillary temperature was set to 275°C, RF lens was set to 40%, precursor ion mass spectrum was acquired at a resolution of 120,000 with a mass range of 380-1400 m/z and a precursor ion charge state range of 3^+^-8^+^. MS^1^ spectra were recorded with a Data Dependent Analysis (DDA) Top20 method, Automatic Gain Control (AGC) was set to a target of 400,000 (100%), maximum injection time mode was set to Auto, precursor ions were isolated with a 1.6 m/z window using the quadrupole mass filter and Monoisotopic Precursor Selection (MIPS) was enabled. For Orbitrap mass analysis based MS^2^ acquisition, stepped higher energy collision induced dissociation (HCD) was applied with a normalized energy of 25, 35%. Orbitrap resolution was set to 30,000, Mass Range was set to Normal, Automatic Gain Control was set to 200% and Maximum Injection Time was set to 54 ms.

Data analysis was performed using Thermo Fisher Scientific Proteome Discoverer 2.5, with the Sequest HT search engine node used to search the RAW files generated from LC-MS/ MS. Default settings were selected unless otherwise stated. RAW files were searched against a bespoke FASTA file containing the QBP sequence and 200 randomly selected protein sequences from *Homo sapiens*. Modifications included dynamic methionine oxidation (+15.995 Da), static cysteine carbamidomethylation (+57.021 Da), dynamic N-terminal acetylation (+42.0106 Da), dynamic lysine DSS-d_0_ amidated monolink (+155.095 Da), and dynamic lysine DSS-d_4_ amidated monolink (+159.120 Da) (note that search for DSS modification is identical to BS3). Monolinks that were specific to the glutamine-free QBP open conformation were obtained by subtracting modification sites that were also found in the closed, glutamine-bound conformation. The remaining monolinks are listed in Table S1, which includes the lysine K:137 that was predicted to be exposed in the open conformation, but occluded in the closed conformation of QBP (annotated MS/MS spectrum in Fig. S3).

### Predicting an ensemble of conformations with AF2 and selecting near-native models with XLP and MP

Conformational ensembles were predicted for luciferase and QBP using a modified version of AlphaFold v2.1.1 (1), that limits the MSA depth, resulting in a greater variation in predicted conformers compared to the default conditions (18). Alignment depths of 8, 16, 32, and 64 were used for template-free AF2 prediction for a total of 200 predicted models (50 per alignment depth).

For C3, which is cleaved into two chains, AlphaFold-Multimer (AF-MM) v2.1.1 (30) was employed to model the complex. In this case, ten conformers were predicted per model, resulting in 50 AF-MM models. Alignment depth was not limited for C3, as the feature was not yet implemented for AF-MM. To avoid bias towards PDB-deposited structures, the max-template-date parameter was set to a date prior to the first deposited related structure, according to PDB’s structural similarity query function (31).

The generated AF2 models were filtered based on their predicted TM (pTM) scores, an internal measure of modeling confidence in AlphaFold. The distributions of pTM scores for each protein in the test dataset can be found in Fig. S4, along with the minimum score threshold used to remove models from further analysis.

Subsequently, XLP and MP scores were computed for each AF2 model and conformation. The AF2 models were ranked by decreasing XLP and MP scores, and the top-scoring models for each protein and conformation were visualized using UCSF ChimeraX (32) .

### Interpolating conformational ensembles for C3 and luciferase

To address the potential impact of limited sampling by AF2, we created structural ensembles for C3 and luciferase by morphing the structures from the open to closed conformation. This was accomplished using the morph function in ChimeraX, which generates intermediate conformations by interpolating along internal coordinates. The resulting ensembles, consisting of 50 models each, were subjected to energy minimization using GROMACS with the AMBER94 force field (33, 34). Subsequently, the XLP score was applied to these morphed ensembles using experimentally determined crosslinks obtained from both the open and closed conformations. The top-scoring models were visualized using ChimeraX for further analysis.

## Results

### Benchmarking monolink and crosslink probability scores with simulated XL-MS data

We first evaluated the performance of our model assessment scores, MP and XLP, using simulated monolink and crosslink data, by measuring the ranking accuracy (Spearman CC of the scores vs RMSD) and near-native selection (AUC) of each score on the 3DRobot dataset.

Figure 3 illustrates the performance of MP and XLP scores, compared to the Matched and Non-accessible Crosslink score (MNXL). All scoring functions demonstrated improved ranking accuracy (CC, Figure 3A) and near-native structure selection (AUC, Figure 3B) with increasing recovery rates. This is expected, as more crosslinks or monolinks provide additional constraints, narrowing down the range of possible conformations. Notably, XLP outperformed MNXL in both ranking accuracy and near-native selection, especially in the low crosslink recovery regime. Removing depth information from XLP, which leaves only Euclidean distance, resulted in decreased performance compared to both XLP and MNXL, indicating the importance of depth information.

**Fig. 3.**
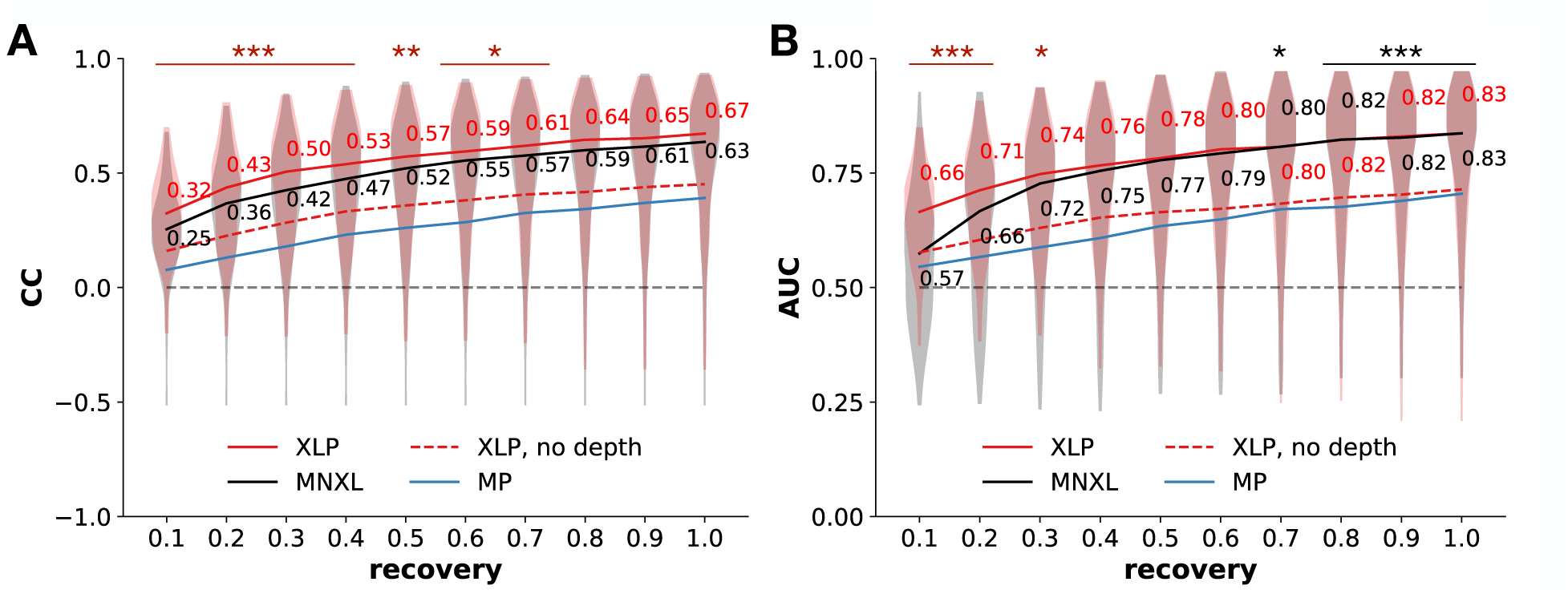
Performance of crosslink and monolink scoring functions in terms of (A) ranking (Spearman CC vs rmsd) and (B) distinguishing near-native structures (AUC) in the 3DRobot dataset across a range of 10-100% crosslink or monolink recovery. The horizontal dashed gray line in each case defines performance that is no better than chance, while the other lines depict the median value for each distribution. Overall, XLP performs best among the scores, particularly at low crosslink recovery rates (≤ 30%). Red asterisks denote recovery rates at which XLP significantly outperforms MNXL, while black asterisks denote the reverse (*** p < 0.001, ** p < 0.01, * p < 0.05). Removing depth information from XLP degrades performance to below MNXL, but above the MP score.

The monolink score MP showed the lowest performance among the assessed scores. However, it still demonstrated near-native selection accuracy comparable to XLP without depth information, particularly at higher recovery rates (Figure 3B).

### Conformational variability predicted by Alphafold2 using shallow MSAs

We employed AlphaFold2 to predict ensembles of models for C3, luciferase, and QBP using shallow multiple sequence alignments (MSAs) as described in the Experimental procedures. Figure 4A displays the resulting AF2 ensembles, illustrating each model’s root-mean-squared deviation (RMSD) from its respective PDB open and closed conformations. Notably, near-native models were observed for all cases except for the closed conformation of C3 and the open conformation of luciferase.

**Fig. 4.**
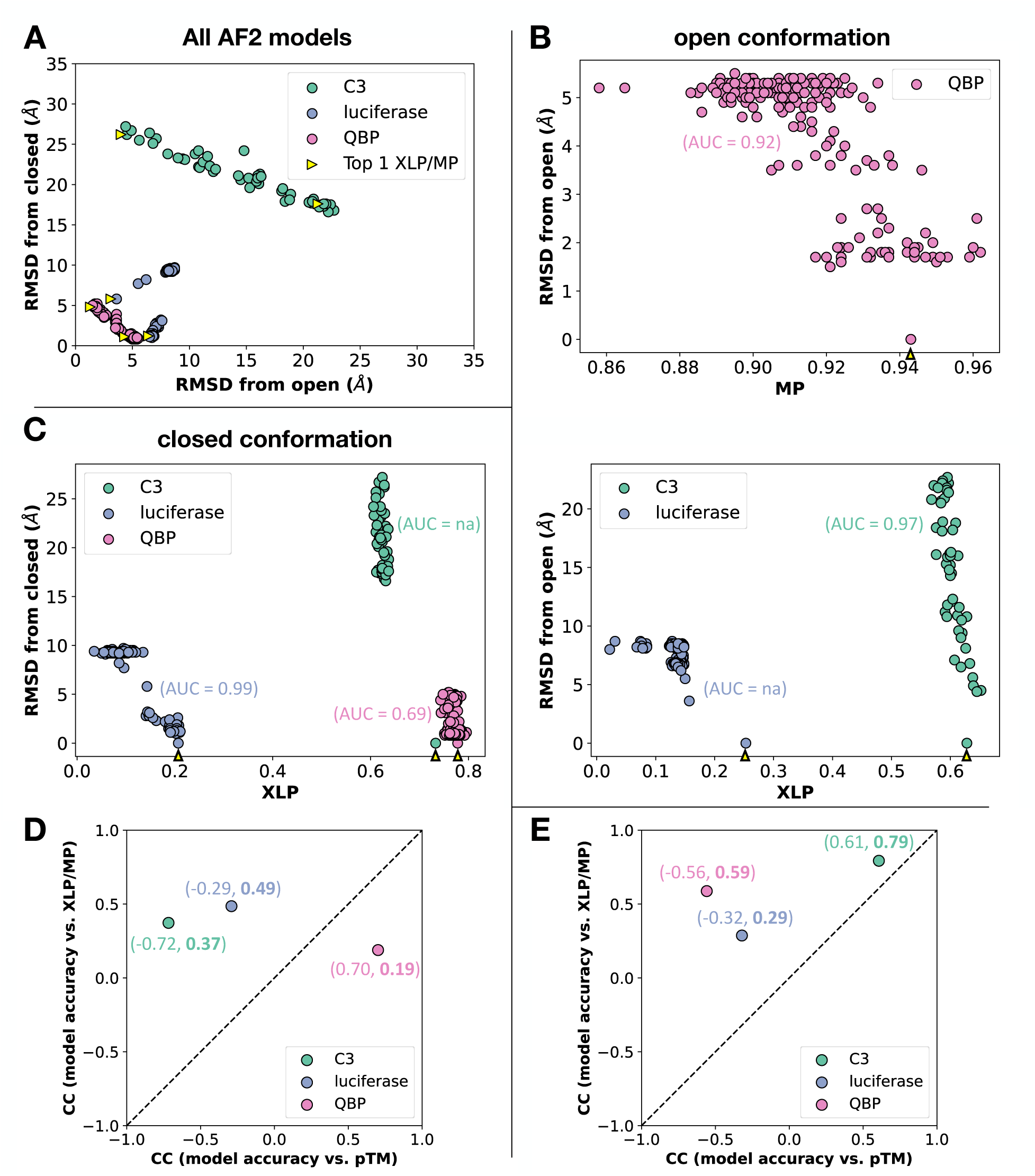
(A) RMSD of AF2 models for C3, luciferase, and QBP from their respective open and closed PDB conformations. Each point represents an AF2 model. Yellow arrows point to the Top 1-XLP/MP scoring models. In all cases, XLP or MP was either able to select a near-native model, or in the absence of near-native structures, to select one of the lower-RMSD structures from the set of AF2 predictions. For C3 and QBP, the AF2 structures lie along a straight line between the open and closed conformations, indicating that the AF2 ensemble might represent physically-relevant intermediate structures between the two conformations. (B) AF2 models with their MP and XLP scores plotted against RMSD from the known PDB open conformation, and (C) against the closed conformation. The PDB conformations are indicated by yellow arrows. The AUC for near-native selection is indicated along each case in parenthesis (na, stands for no near-native structures found). (D - E) Ranking accuracy comparisons between XLP/MP and pTM show that the XL-MS based scores performed better (above the diagonal) than pTM for all cases in closed (D) and open (E) conformations, except QBP closed conformation.

An interesting observation was that the models for C3 and QBP exhibited a linear distribution between the open and closed conformations. This suggests that these ensembles encompass a range of conformations, including near-open and near-closed states, as well as intermediate configurations between these “endpoints”. However, for luciferase, the majority of model variation occurred away from the open-to-closed axis, albeit closer to the closed conformation. This indicates the presence of potentially misfolded structures. This observation is further supported by the 3D plot in Figure S5, which demonstrates relatively low pTM scores off this axis for luciferase and QBP, indicating that AF2 is less confident about these off-axis models.

### Filtering conformational variability from AF2 with XLP and MP

We calculated XLP and MP scores for each AF2 model using XL-MS data from either the open or closed conformation. In all cases, the model with the highest XLP or MP score corresponded to a near-native structure, or, if a near-native model was not available, to a conformation within 1 Å of the most accurate model in the ensemble (Figure 5). The near-native selection accuracy, as measured by the AUC, ranged from 0.69 for the closed conformation of QBP to ≥0.92 for the other cases where AF2 predicted a near-native model (Figure 4B and C). In contrast, when using the pTM score, the AUC varied across different conformations, showing selective behavior for the open conformation of C3 and the closed conformation of QBP (AUC ∼ 0.9), anti-predictive behavior for the open conformation of QBP (AUC = 0.31), and near-random performance for the closed conformation of luciferase (AUC = 0.47) (Fig. S6). XLP and MP scores also exhibited more accurate ranking of models based on RMSD compared to pTM, except for the closed conformation of QBP (Figure 4D and E). These findings demonstrate that the XLP and MP scores enable the identification of the conformer represented by a given set of XL-MS data from an ensemble of structural models. In contrast, the use of pTM alone is, at best, only selective for one conformation.

**Fig. 5.**
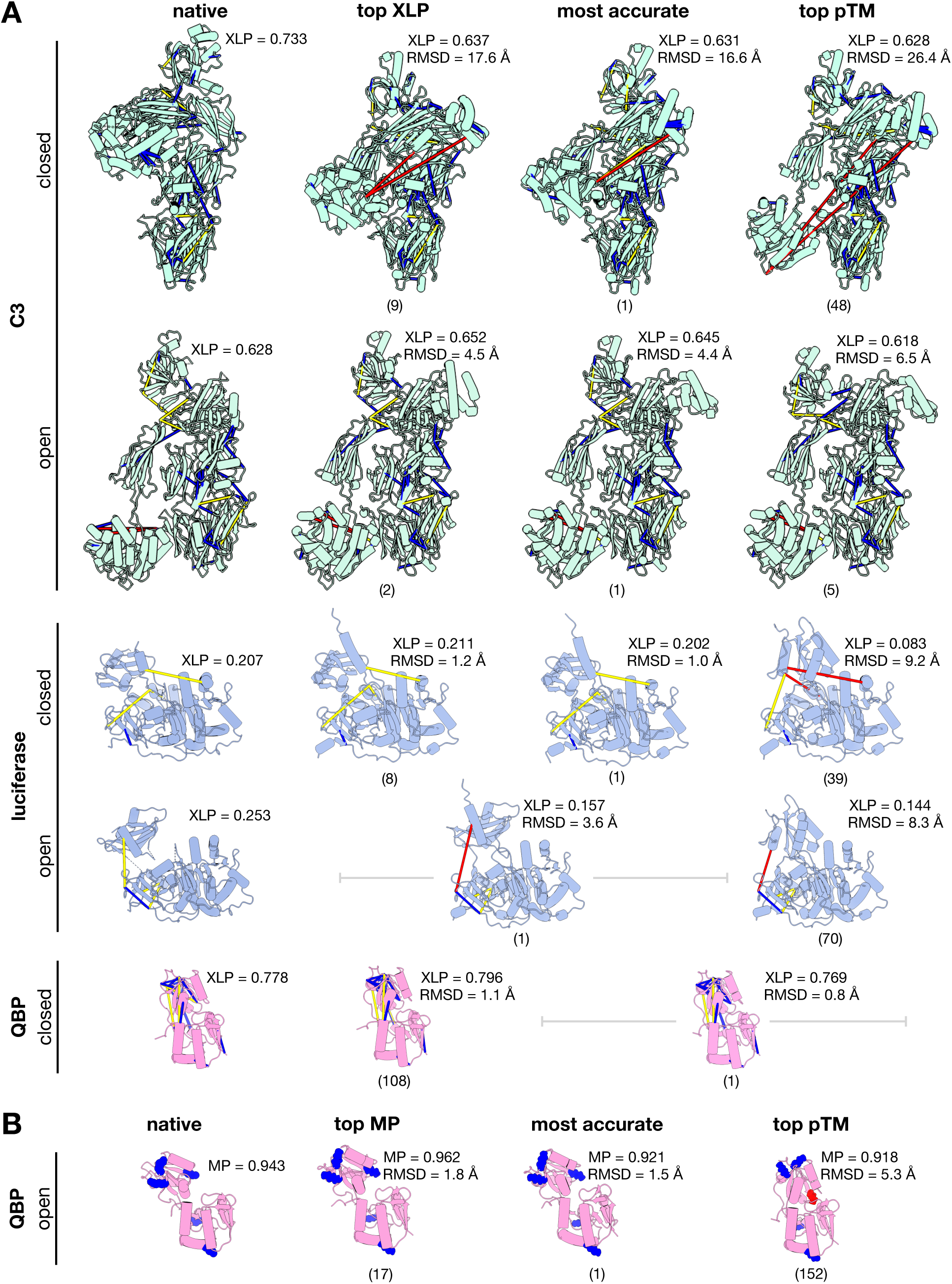
AF2 models and the PDB structures of open and closed conformations of C3, luciferase and QBP (A) with the crosslinks shown, colored by Cα-Cα distance (blue: < 21Å, yellow: 21-33Å, red > 33Å), and (B) with the monolinks shown, colored by residue depth (blue: 6.25Å, red: >6.25Å). Three representative structures are shown for each AF2 ensemble: the top XLP or MP scorer, the most accurate (lowest RMSD from PDB structure), and the model with the highest pTM score. The corresponding XLP or MP scores and RMSD from the target conformation are shown alongside each model, with the rank of the model in terms of accuracy shown below each structure in brackets.

It is self-evident, however, that scoring functions cannot select for the correct answer if it is not among the options. To address this issue in the case of C3 and luciferase, we disentangled the effect of poor AF2 sampling by building an interpolated ensemble consisting of the open and closed conformation, and intermediate structures between them (Fig. S7A). Using these interpolated ensembles resulted in very good performance of XLP in ranking (CC > 0.89), and near-native selection (AUC > 0.95) for C3 open and closed, and luciferase open conformations, and reasonable ranking and near-native selection for luciferase closed conformation (CC = 0.68, AUC = 0.73) (Fig. S7B). For all cases, the top-XLP scoring models were near-native, and within the top-10 models in terms of RMSD to the target (Fig. S7C).

## Discussion

AlphaFold2 represents a paradigm shift in structural biology in many ways, but it does not preclude the use of experimental methods, particularly in the case of flexible proteins. We show here that AF2 can fit very well in an integrative modeling pipeline for flexible proteins by providing an ensemble of structures that can be evaluated using distance constraints and solvent accessibility information from cross-linking mass spectrometry.

In developing scoring functions based on XL-MS data, we wanted one that was more nuanced than just a count of maximum distance violations (9–12), to rank structures that are all possible within the relatively-large distance constraint imposed by BS3 and DSS (10, 16). Within the umbrella of existing crosslink-based scoring functions, those based on solvent accessible surface distance (SASD) were found to perform better than Euclidean distance (ED) (14–17). However, SASD computation is more time-consuming, as it involves tracing the possible paths of the crosslinker along the protein surface, whereas ED only requires a single distance computation. We show here that the performance of the SASD-based score MNXL can be approximated, and even exceeded, by our ED-based scoring function XLP (crosslink probability), thereby circumventing the need for more time-consuming SASD computations, and opening up the possibility of using XLP to assess large conformational ensembles. These results are consistent with our previous findings that show that including solvent accessibility can improve the performance of an ED-based scoring function to almost the level of the SASD-based scoring function MNXL (16, 35). Instead of solvent accessibility, in our new scoring function we used residue depth, resulting in improved performance of XLP over MNXL, notably at low crosslink recovery rates of 30% or less. Performance improvements in the low crosslink recovery regime are particularly impactful, since experimental crosslink recovery rates are typically reported to be in this range (14, 16, 35–37), particularly in the noisier conditions found in in-situ crosslinking (38, 39). The XLP score was also shown to distinguish between different conformations of the proteins in our test dataset, with the top scoring model in each case either being the most accurate one, or within 1 Å of the most accurate model.

We also conceptualized and benchmarked a scoring function based on monolinks, the monolink probability (MP) score, which is a measure of the likelihood that an amino acid at a certain depth from the protein surface will be tagged with the crosslink reagent. Previously, we have demonstrated that although monolinks contain less information than crosslinks, they can also be used to select for near-native structures from a set of decoys, albeit at higher recovery rates of at least 50% (24). Our current work has confirmed these previous results. Furthermore, the MP score was needed to select for the open conformation of QBP as there were no crosslinks found exclusive to this conformation. This is due to the small size of QBP, such that all possible crosslinking events can be spanned by BS3/DSS whether in open or closed conformation. What was informative in this case is the reduced residue depth that occurs when previously inaccessible regions of the protein are exposed to solvent when it is in the open conformation. Surprisingly, even one differentially-exposed lysine residue was sufficient to distinguish the open conformation of QBP from the rest of the conformational ensemble (Fig. 4B and 5B), indicating the potential of this approach in cases where more numerous conformation-specific monolinks could be found. Differential residue depth data can be obtained from experiments such as hydrogen-deuterium exchange mass spectrometry (HDX-MS) (40), but one can also get this “for free” in XL-MS experiments in the form of monolinks, which are currently not frequently used for data analysis. We hope this demonstration of the importance of monolinks will encourage experimentalists to use monolink data more often, and to deposit them in publicly-available databases such as the Proteomics Identifications Database, PRIDE (41).

Aside from the lack of deposited monolink data, we are limited by the current state of structure prediction software. In this case, running AF2 with a shallow MSA allowed the endpoint PDB structures of QBP to be predicted, as well as plausible intermediate structures. This is less the case for C3, for which only models near the open conformation were predicted, as well as some putative intermediate structures, but no conformations at or near the PDB closed conformation. For luciferase, the structural variation captured by AF2 occurs along a completely different axis than the endpoint structures documented in the PDB, suggesting that these predicted conformations might not be physically-relevant. This tradeoff between modeling accuracy and conformational variability as a function of alignment-depth has been discussed in greater depth in the publication detailing the technique (18).

We have also observed, during both the benchmarking and testing of XLP, that crosslink recovery is directly proportional to the ranking accuracy and near-native selection. It has also been previously shown that the location of crosslinkable sites in the protein also limits their potential information content (35, 42). For example, if both ends of a crosslink span a rigid region of an otherwise flexible protein, then it would not distinguish between different conformations of the protein. This position dependence can be ameliorated by increasing crosslink recovery with the use of crosslinking reagents with different reactive end groups to increase the likelihood of getting informative crosslinks in an XL-MS experiment. Another way to increase the discerning power of XL-MS based scoring functions is to use shorter crosslinkers, thereby providing narrower ranges of distance constraints (43). This is particularly important to resolve the differences in conformation in a small protein, like QBP in our dataset.

It should be noted, however, that since our proposed scoring function is based on empirical distributions of spanned C⍺-C⍺ distances, this distribution would have to be computed separately for different crosslinking reagents, thus further highlighting the need to deposit experimental XL-MS data in PRIDE.

## Acknowledgments

This work was supported by Universität Hamburg and Hamburg-X grant LFF-HHX-03 to the Center for Data and Computing in Natural Sciences (CDCS) from the Hamburg Ministry of Science, Research, Equalities and Districts (BWFGB), the Leibniz Institute of Virology as part of Leibniz ScienceCampus InterACt (funded by the BWFGB Hamburg and the Leibniz Association) (to MT and KG), a Wellcome Collaborative Award in Science (209250/Z/17/Z) (to KG, KT and MT), and a Wellcome Trust Multiuser Equipment grant (221521/Z/20/Z) (to KT).

## Notes

### Competing Interest Statement

The authors have declared no competing interest.

