## Supplementary Information for "Modeling flexible protein structure with AlphaFold2 and cross-linking mass spectrometry"

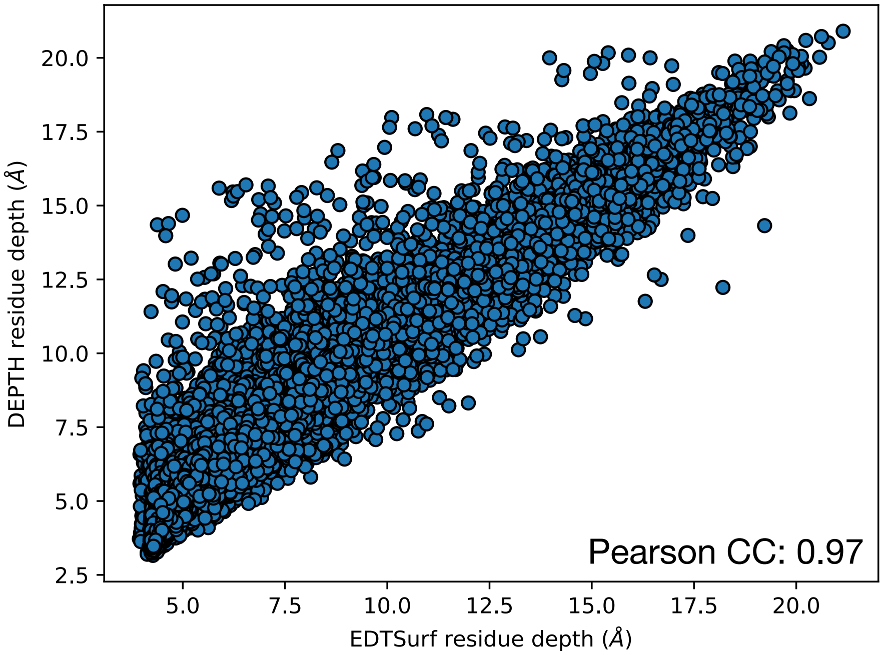


**Fig. S1**. Computed depths of lysine residues from BS3/DSS-tagged proteins in the XlinkAnalyzer database. Residue depths were computed using a computing-intensive explicit solvent method, DEPTH (22), and a faster method using Euclidean distance transform (EDTSurf) (20, 21). EDTSurf can approximate the residue depths computed by DEPTH, with a Pearson correlation coefficient of 0.97.

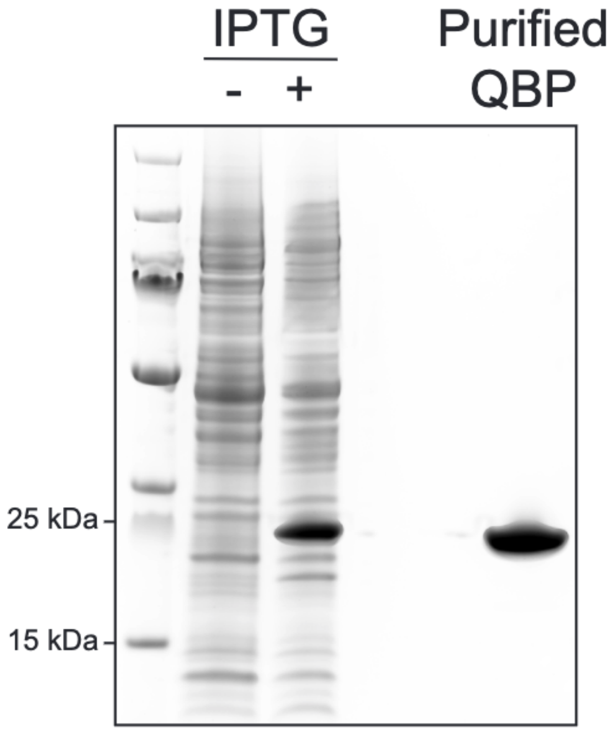


**Fig. S2.** Purification of QBP. Results of Coomassie stained SDS-PAGE showing whole cell lysates of *E.coli* before (-) and after induction of QBP expression by 500 µM IPTG for 5 h at 30°C (+). Amounts of whole cell lysates were normalized to measured $OD_{600}$ prior to loading. Purity of QBP preparation after Ni-sepharose affinity and size exclusion chromatography is demonstrated by loading of 8 µg purified QBP.

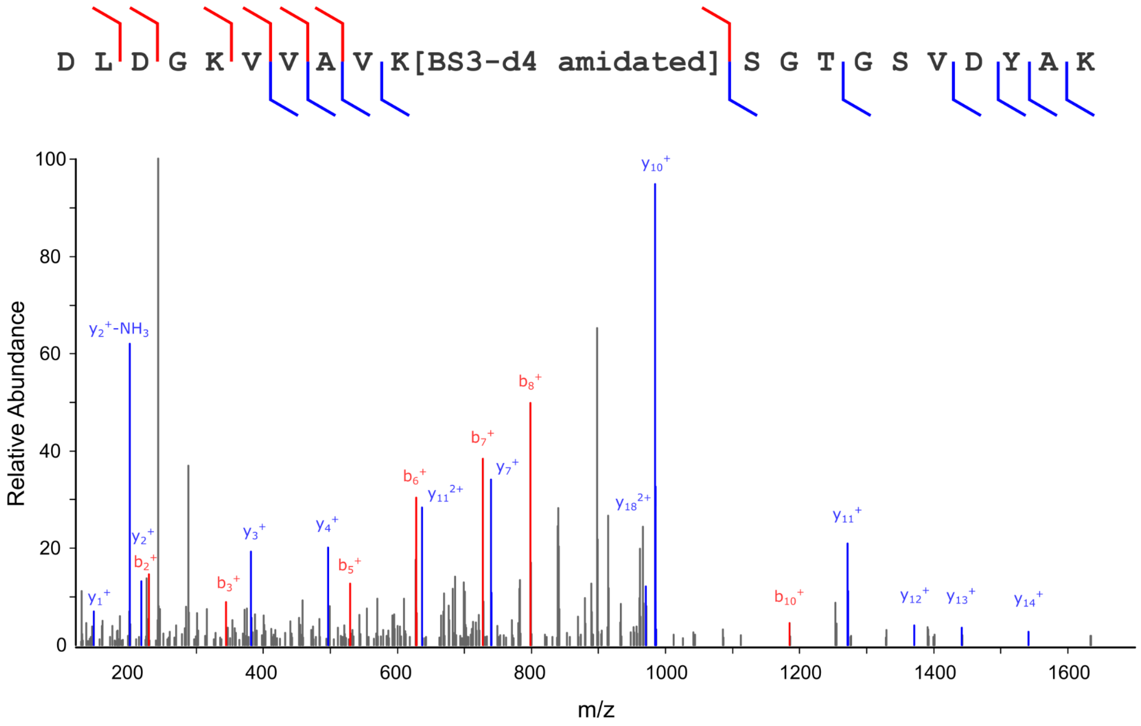


**Fig. S3.** MS/MS spectrum from higher energy collision induced dissociation of precursor ion: [DLDGKVVAVK(BS3-$d_4$ amidated)SGTGSVDYAK]$^{3+}$, corresponding to lysine-137 monolink identification. Precursor ion mass [M+H]$^+$: 2168.17691, *m/z*: 723.39717. Blue peaks represent *y* ions, red peaks represent *b* ions, with fragmentation sites annotated across the peptide sequence.

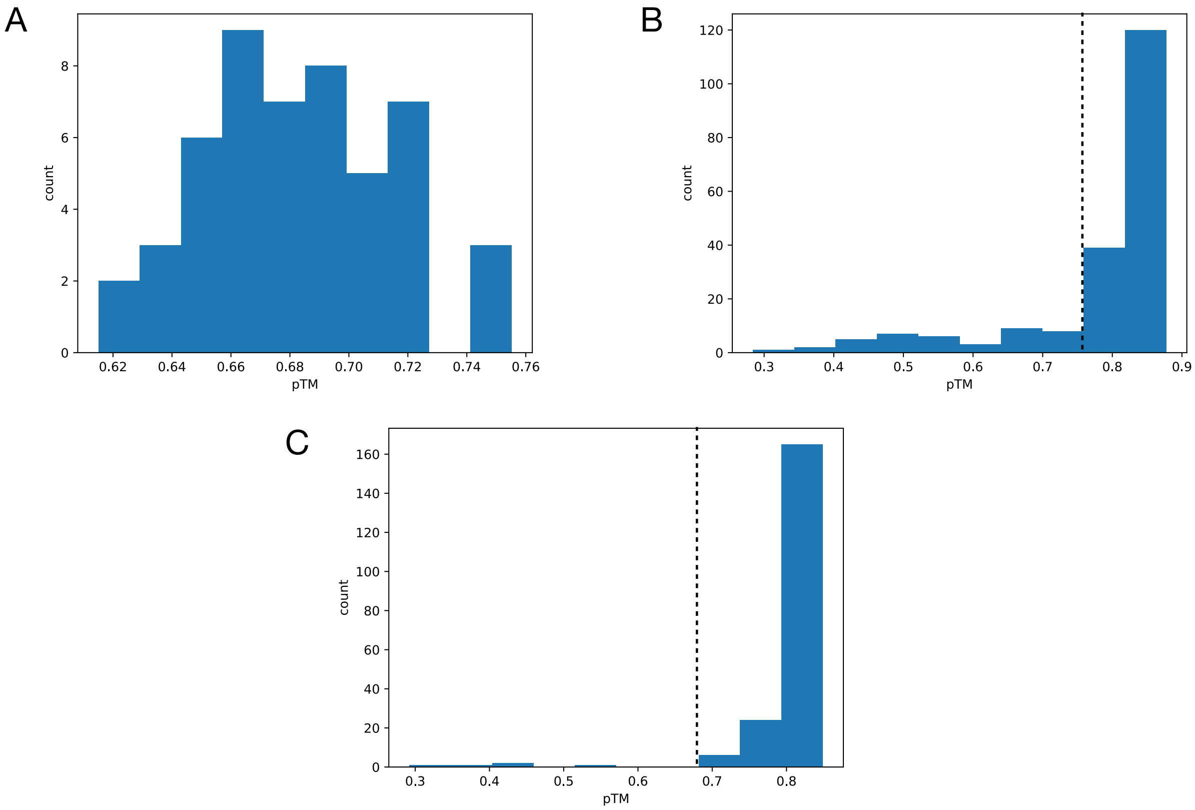


**Fig. S4.** Distributions of predicted TM-scores (pTM) for AF2 models of (A) C3, (B) luciferase and (C) QBP. Models with low pTM values were excluded by removing models below the pTM threshold, which are indicated by the dashed lines.

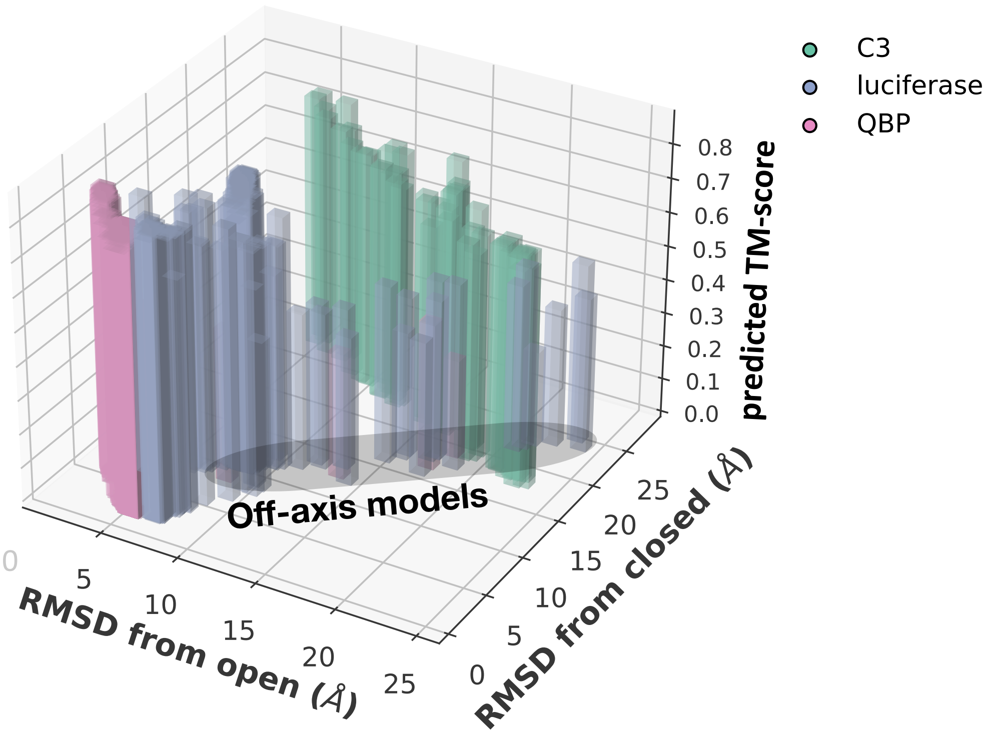


**Fig. S5.** Spread of AF2 models of C3, luciferase and QBP with respect to RMSD against the PDB open and closed conformations, along with the predicted TM-score of each model. For C3 and QBP, models were predicted that were either relatively near the open or closed conformation, or somewhere between the two conformations. For luciferase and QBP, models that are off this open-to-closed axis have notably lower pTM scores, indicating potentially misfolded structures.

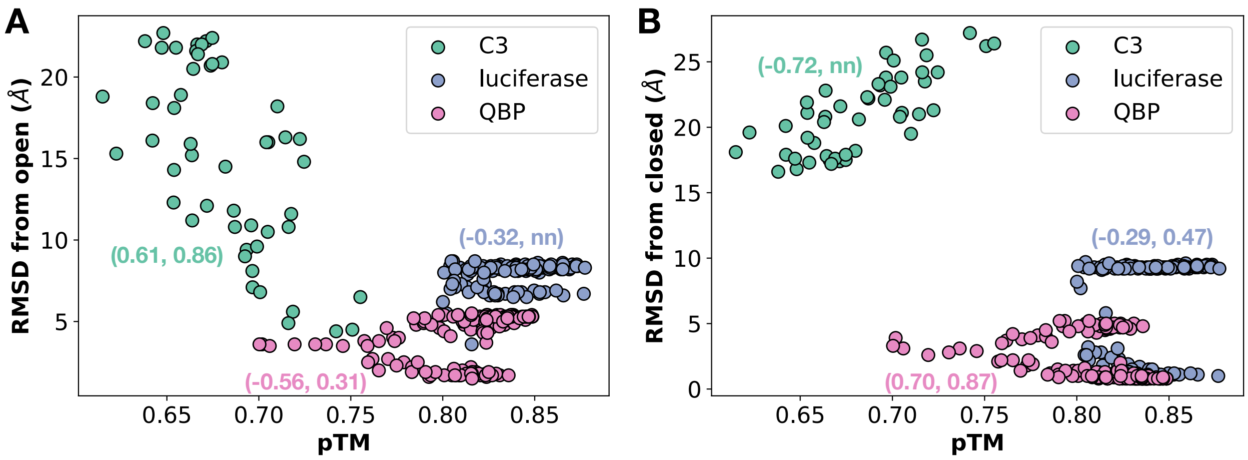


**Fig. S6.** RMSD of AF2 models of C3, luciferase and QBP from the PDB open (A) and closed conformations (B), as a function of predicted TM-score. The tuples in parenthesis are the (Spearman CC vs RMSD, AUC), which are measures of ranking accuracy and near-native selection of the pTM score, respectively. Higher pTM scores select for models near the C3 open conformation, and QBP closed conformation, but are not selective for other conformations

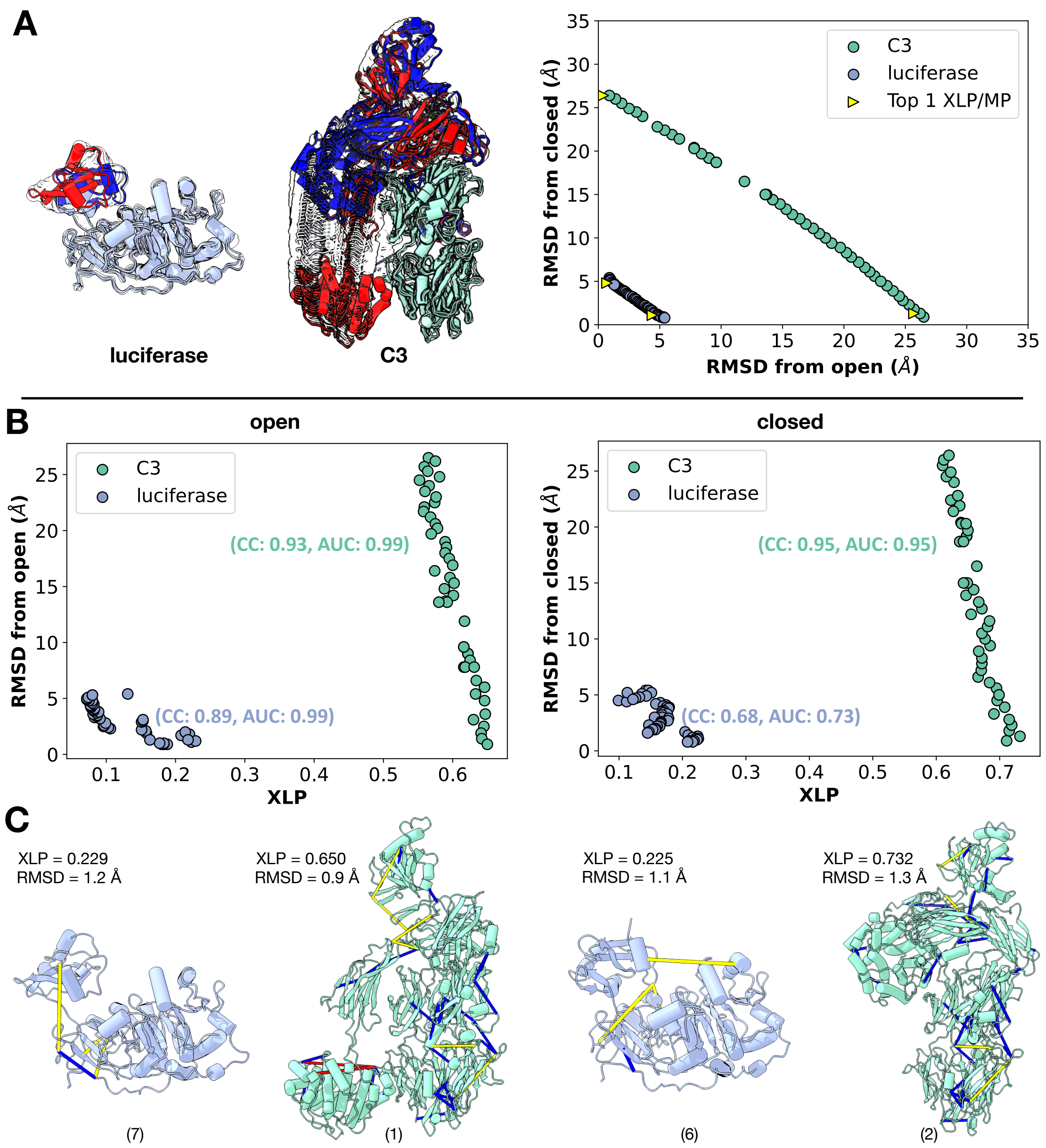


**Fig. S7.** (A) Ensembles of luciferase and C3, obtained by interpolation between PDB open and closed conformations. In both cases, the mobile regions of the open conformation are colored red, while the closed conformation is shown in blue. The interpolated ensemble contains models that are near-open and near-closed, as well as intermediate structures between the two (RMSD plot, right) (B) The XLP score ranked models of C3 and luciferase by correctness (CC > 0.68), as well as distinguished near-native structures from decoys with reasonable accuracy (AUC > 0.70). (C) The top-XLP scoring models in the interpolated ensemble. Below each model is its rank in terms of RMSD to the native structure.

**Table S1. Experimentally-determined crosslinks and monolinks used for XLP/MP computation**

| Protein | Conformation: open | Conformation: closed |
|---|---|---|
| **C3** | B:232-B:943, B:369-B:379, B:825-B:917, A:267-B:759, A:44-B:369, A:44-B:378, A:44-B:531, A:44-B:545, A:1-A:464, B:433-B:499, B:441-B:467, B:441-B:499, B:496-B:508, B:531-B:572, B:634-B:653, B:879-B:927, B:927-B:943, A:183-A:593, A:219-A:406, A:219-A:585, A:242-B:219, A:331-A:600, A:337-A:600, A:340-A:611, A:343-A:406, A:343-A:611, A:363-A:406, A:396-A:609, A:544-A:562, A:562-B:111, A:585-A:593, A:586-A:593, A:586-A:600, B:189-B:245, B:258-B:653, B:258-B:681, B:287-B:634, A:227-A:267, A:396-A:611, B:681-B:829 (15% recovery) | B:399-B:759, B:696-B:825, B:829-B:923, B:20-B:399, B:255-B:764, A:578-A:588, B:10-B:49, B:10-B:50, B:13-B:50, B:14-B:49, B:16-B:20, A:1-A:464, B:433-B:499, B:496-B:508, B:531-B:572, B:634-B:653, B:701-B:709, B:854-B:863, B:879-B:927, B:927-B:943, A:183-A:593, A:219-A:406, A:219-A:585, A:242-B:219, A:331-A:600, A:337-A:600, A:340-A:611, A:343-A:406, A:343-A:611, A:363-A:406, A:396-A:609, A:544-A:562, A:562-B:111, A:585-A:593, A:586-A:593, A:586-A:600, B:189-B:245, B:258-B:653, B:258-B:681, B:287-B:634, A:227-A:267, A:396-A:611, B:696-B:709, B:258-B:764 (16% recovery) |
| **Luciferase** | A:380-A:541, A:9-A:380, A:9-A:329, A:372-A:329 (4% recovery) | A:380-A:529, A:303-A:529, A:142-A:534, A:9-A:372 (4% recovery) |
| **QBP** | A:98, A:117, **A:137**, A:196, A:221 (100% recovery of informative lysine crosslink, in bold) | A:26-A:68, A:62-A:221, A:98-A:213, A:99-A:213, A:109-A:227, A:109-A:240, A:213-A:221, A:117-A:127, A:117-A:132, A:117-A:137, A:124-A:132, A:124-A:137, A:124-A:196, A:127-A:137, A:127-A:147, A:127-A:151, A:127-A:153, A:132-A:151, A:132-A:153, A:147-A:153, A:98-A:147, A:99-A:147, A:109-A:127, A:117-A:241, A:188-A:240, A:188-A:241 (25% recovery) |
